# Micro-XRF study of the troodontid dinosaur *Jianianhualong tengi* reveals new biological and taphonomical signals

**DOI:** 10.1101/2020.09.07.285833

**Authors:** Jinhua Li, Rui Pei, Fangfang Teng, Hao Qiu, Roald Tagle, Qiqi Yan, Qiang Wang, Xuelei Chu, Xing Xu

## Abstract

*Jianianhualong tengi* is a key taxon for understanding the evolution of pennaceous feathers as well as troodontid theropods, and it is known by only the holotype, which was recovered from the Lower Cretaceous Yixian Formation of western Liaoning, China. Here, we carried out a large-area micro-X-Ray fluorescence (micro-XRF) analysis on the holotypic specimen of *Jianianhualong tengi* via a Brucker M6 Jetstream mobile XRF scanner. The elemental distribution measurements of the specimen show an enrichment of typical bones couponing elements such as S, P and Ca allowing to visualize the fossil structure. Additionally, to this, the bones are enriched in several heavier elements such as Sr, Th, Y and Ce over the surrounding rocks. The enrichment is most likely associated to secondary mineralization and the phosphates from the bones. Interestingly the plumage shape correlates with an enrichment in elements such as Cu, Ni and Ti, consistent with a previous study [1] on *Archaeopteryx* using synchrotron imaging. The analysis presented here provide new biological and taphonomic information of this fossil. An *in-situ* and nondestructive micro-XRF analysis is currently the most ideal way to map the chemistry of fossils, so far this is manly restricted to small samples. Larger samples usually required a synchrotron facility for analysis. Our study demonstrated that laboratory-based large-area micro-XRF scanner can provides a practical tool for the study of large large-sized specimens allowing collect full chemical data for a better understanding of evolutionary and taphonomic processes.

## I. Introduction

Morphological studies on the skeleton and the plumage have forged a solid link between non-avialan coelurosaurians and birds building a well-accepted framework to understand the dinosaur-bird transition [2–4]. Recent works started to look beneath the surface and into the ultrastructure and chemistry in fossil feathers and bones and bring our understanding of this major transition to another new level [1, 5–12]. Chemistry of exquisitely preserved fossil animals including several iconic flying/gliding capable theropods have been investigated to reveal information of their paleobiology and the fossilization process [1, 6, 9, 10, 12–14]. Various chemical imaging techniques, e.g., Fourier-transform infrared (FTIR) spectroscopy, Raman spectroscopy, X-ray spectroscopy, and secondary-ion mass spectroscopy (SIMS)), have been employed to track the molecular, elemental and isotopic information [1, 5, 6, 8–13, 15–18]. Unfortunately, previous studies all focused on non-destructive chemical imaging on small-sized specimens (e.g., *Archaeopteryx, Confuciusornis*) as limited by the analyzing instruments [6, 8, 11, 12], or chemical analyses on small samples destructively taken from fossils [9, 10, 13, 16, 17]. So far systematic non-destructive chemical imaging on large-sized specimens of non-avialan dinosaurs have not been done.

The Middle-Upper Jurassic and Lower Cretaceous of western Liaoning and neighboring areas have yielded abundant avialans and non-avialan dinosaur specimens with exquisite preservation of the soft tissues that enlightened us the origin and early evolution of feathers and bird flight [3, 19–22]. The unique preservation of these specimens also provides a good opportunity to detect the hidden ultrastructural, chemical and taphonomic information of feathers and bones [8–10, 16, 23]. The troodontid theropod *Jianianhualong tengi* is represented by an exquisitely preserved specimen from the Lower Cretaceous Yixian Formation [24]. It has been revealed to bear asymmetrical pennaceous feathers using regular tools and Laser-stimulated fluorescence imaging, suggesting feather asymmetry a synapomorphy of the Paraves, a clade including birds and their close relatives. Compared to other Mesozoic paravians with asymmetrical pennaceous feathers, *Jianianhualong* is relatively large and apparently not adapted for flight as indicated by its short forelimbs [24]. Besides the novelty on feathers, *Jianianhualong* is also featured by a mosaic combination of primitive and derived troodontid features [24].

Here we performed several micro-X-Ray fluorescence analysis (micro-XRF) on the holotype of *Jianianhualong tengi* with different spatial resolution to reveal the chemistry of the whole specimen as well as some relevant features. The analysis provides new information for understanding the paleobiology of this troodontid dinosaur and the taphonomy of this important fossil.

## 2. Materials and methods

### 2.1 Troodontid dinosaur *Jianianhualong*

The holotype, the only known specimen of *Jianianhualong tengi*, was discovered from the Lower Cretaceous Yixian Formation (the middle section of the Jehol group) of Baicai Gou, Yixian County, western Liaoning, China. The holotype (DLXH 1218) is now housed at the Dalian Xinghai Museum (DLXH). Unlike other troodontid specimens from the Yixian Formation that are 3D-preserved in tuffaceous sandstones, DLXH 1218 is compressed and preserved on a tuffaceous shale slab, even though the bones are not entirely flattened (Fig. 1a). DLXH 1218 contains a nearly complete skeleton associated with soft tissues including feathers (Fig. 1a). The skeleton is black in color. At least two layers of the matrix can be observed on the slab: an overlying white layer (WL) and an underlying brown layer (BL, possibly rich in organic matters) that preserve the fossil. The regions immediate surrounding the skeleton, possibly soft remains (SR) of the specimen, have brighter color than the rest parts of the matrix of the brown layer (Fig. 1a). The size of the slab is approximately 90 cm in length and 70 cm in width. DLXH 1218 measures ~100 cm in preserved skeletal body length and ~117 mm in femoral length. DLXH 1218 is estimated to be ~112 cm in total length with a fully preserved tail.

**Fig. 1.**
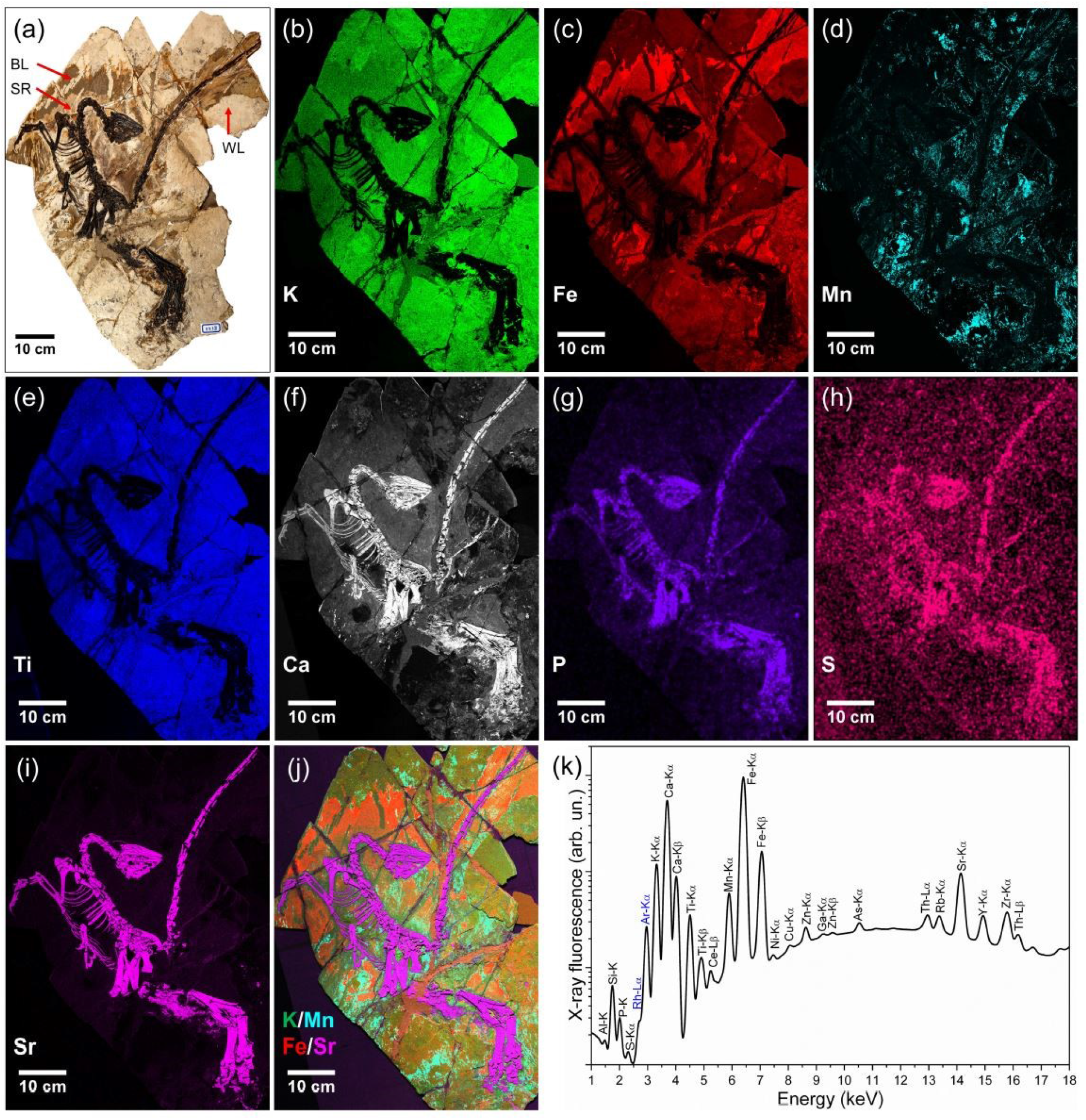
Overall element distribution of the holotype of *Jianianhualong* revealed via micro-XRF imaging. (a) Light photo of *Jianianhualong* (DLXH 1218). The skeleton of fossil is represented by the black color. The matrix contains an overlaying white layer (WL), a brown layer (BL), and an immediate layer surrounding the skeleton, possibly soft remains (SR) of the specimen. (b)-(i) Intensity maps in the rays of K-Kα (b), Fe-Kα (c), Mn-Kα (d), Ti-Kα (e), Ca-Kα (f), P-K (g), S-Kα (h), and Sr-Kα (i). More intensity maps of other elements are shown in the supplementary material (Fig. S1). (j) Combined intensity map of four elements. Green, K; aqua, Mn; red, Fe; magenta, Sr. (k) The corresponding XRF spectrum shows the detectable elements of Al, Si, P, S, K’ Ca, Ti, Ce, Mn, Fe, Ni, Cu, Zn, Ga, Zn, As, Th, Rb, Sr, Y and Zr. The Rh and Ar peaks originate from the Rh-target microfocus-X-ray tube and ambient air, respectively.

### 2.2 Micro-XRF imaging and data analysis

The elemental maps of the specimen DLXH 1218, were obtained using the Bruker M6 Jetstream mobile X-ray fluorescence (XRF) scanner. The analysis was performed at the Dalian Xinghai Museum for *in-situ* studies (Fig. S1a). The Bruker M6 Jetstream consists of a mobile measuring head that can scan the surface of a sample with a XY-motorized stage (Fig. S1b). The maximum range of the XY-motorized stage is of 800 mm × 600 mm (H × W). Mounted on the measuring head is a Rh-target X-ray tube (30 W maximum power, usually operated at 50 kV and a maximum current 0.6 mA). The X-ray beam is guided to the sample through a polycapillary focusing optics. The beam diameter can be regulated by adjusting the distance between the X-ray source and the sample surface, standard settings are between 100 μm and 500 μm. The X-ray signal from the sample are collected by a 30 mm^2^ XFlash silicon drift detector with an energy resolution <145 eV at Mn-Kα. The instrument is equipped with two magnifying optical cameras to document the area of analysis. The mobility of the instrument allows to bring the instrument to the object and to measure the samples directly on-site, reducing risk of a transport damage of valuable samples. This provides ideal conditions for an *in-situ* micro-XRF measurement for high performance element distribution analysis of large samples with high spatial resolution in the 100 μm range [25–27].

Being placed horizontally on a table and properly oriented (Fig. S1b), the specimen DLXH 1218 was scanned fully (the whole specimen) and partially (several sub-areas of interest with higher resolution). The spectra were collected, deconvoluted and examined with the Bruker M6 Jetstream software package. Chemical elements were identified in the scan by examining the sum spectrum and maximum pixel spectra (Bright and Newbury, 2004). The net intensity (cps) of each element was also calculated (Tables S1-S4). In order to obtain XRF images with high contrast, principal component analysis (PCA) was applied to limit the principal and major components of the objects, and then an XRF spectrum for each component was recreated [28].

## 3. Results

### 3.1 Overall element distribution revealed via full-area XRF imaging

Full-area XRF elemental maps of the specimen DLXH 1218 are shown in Fig. 1 and Fig. S2. The acquisition conditions where 1500 × 1090 pixel for a total of 1.6 million pixel/spectra, acquisition time per spectrum was 15 ms and the pixel size 500 μm. The total measurement time was 9h30min.

The XRF measurement provides an overall distribution for at least 20 recognizable elements on the specimen (Fig. 1k). Element net intensities on various spots or regions of interest (ROI) of the specimen are shown in Tables S1-S4. Sorting of element by their main phase, shows that the light rock where the fossil is embedded, is rich in Si, K, Ca, Ti, Mn, and Fe (note that all elements are most likely present as oxides, but the oxygen can’t be detected here). The elements present and their respective intensities, suggest a clay rich sedimentary rock with a carbonate fraction.

The areas of high Mn are scattered on the white layer (represented by the black spots under normal light). This is most likely related to pyrolusite mineralization (MnO_2_). As further quantification is not possible under the given analytical conditions e.g., working distance, orientation of the sample to the head and calibration, the acquire data would be discussed semi quantitatively. The darker parts in the visual image of specimen DLXH 1218, some of them associated to feathers regions, correlate with an increase in Fe and As. The various rocks fragments of the fossil are fixed together by a cement like material, containing some traces of Zn.

Zink is distributed primarily in the white layer of the matrix and the cement like material (Fig. S2i). The Zn level is low in the skeleton and the soft tissue including feather remnants, which contradicts the result observed in the Thermopolis *Archaeopteryx*, where the bone materials have higher Zn level than in the feathers and the matrix [12]. As a contrast, the coliiformes bird has higher Zn level in the feathers, while the matrix and the bone contain less Zn [29].

The bone is, as expected rich in Ca and P, corresponding mineralogically to apatite. The apatite mineral appears strongly enriched several incompatible elements such as: Th, Sr, Y, Zr as well as some REE mainly Ce (Fig. S1, Tables S1-S4). High Sr level associated with other fossil materials was also noticed by Gueriau and co-workers [30].

The regions where feather remains can be observed show an enrichment and correlation pattern of several elements including Mn, Ti, Ni and Cu. This best visible especially the feathers region of the detail scan of the tail see later discussion (Fig. 1d and Fig 2e). Whereas an integration of transitional elements such as Mn, Cu and Ni into the organic component appears more obvious, is the enrichment of Ti more difficult to explain, especially as the correlation with of the feather structure with Ti is even much clear than for the other elements as can be seen in the scan from the tail.

**Fig. 2.**
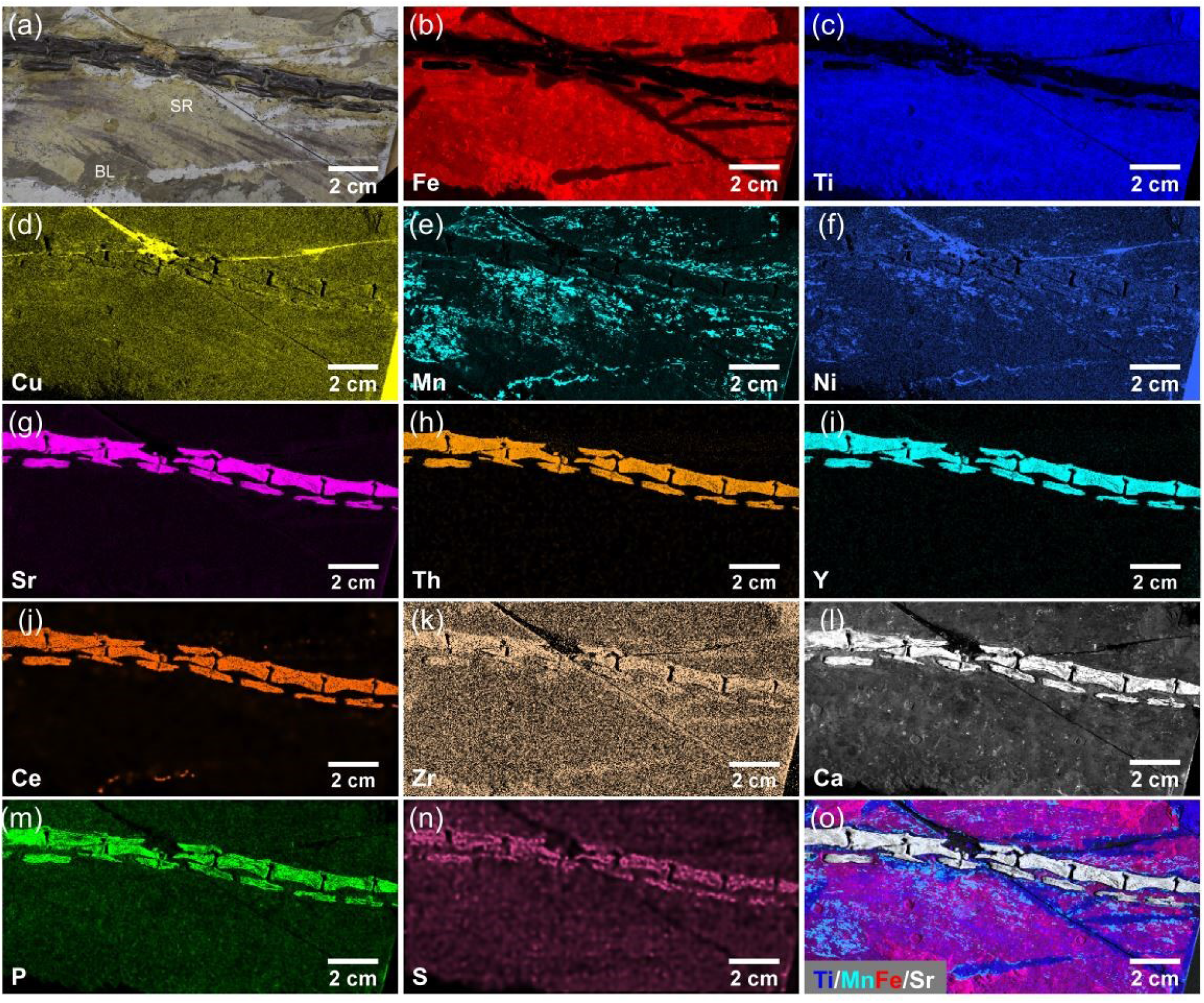
(a) Light photo of a small region of the *Jianianhualong* tail. The bone materials (represented by the black color) and soft tissue (SR) remnants including the feathers, as well as the brown layer (BL) can be observed based on light photo imaging. (b)-(n) Micro-XRF detail maps of Fe (b), Ti (c), Cu (d), Mn (e), Ni (f), Sr (g), Th (h), Y (i), Zr (j), Ce (k), Ca (l), P (m), and S (n) distribution within bone materials and feathers of the *Jianianhualong* tail. (o) Combined intensity map of four elements. Blue, Ti; aqua, Mn; red, Fe; white, Sr.

Rubidium and Ga are possibly controlled by the matrix. They are high in the white layer of the matrix, slightly lower in the brown layer of the matrix, and extremely low in the skeleton (Fig. S2d, S2e).

Iron and As levels are very low in the skeleton. They are most concentrated in the brown layer, and less concentrated in the overlying white layer (Fig. 1c, Fig. S2f). The soft tissue remnants including the feathers (represented by the light brown areas close to the skeleton under visible light) exhibit lower Fe level than their immediate surrounding matrix (the brown layer) (Fig. 1c).

Copper and Ni are distributed in the white layer of the matrix and slightly more concentrated in the feathers. They are less accumulated in the bone materials and the brown layer of the matrix, and high concentrations of both elements appear in the glue which is artificially added during the fossil restoration (Fig. S2g, S2h). A high level of Cu and Ni is also known in the feathers of many fossil and living taxa such as *Anchiornis, Archaeopteryx, Gansus* and other fossil and living birds [1, 6, 8, 12, 29, 31]. In all these specimens and the specimen DLXH 1218, the Cu level is generally higher in the feathers than in the bones, and the copper concentration usually varies on different feather samples of an individual [6, 12, 29, 31].

### 3.2 Elements associated with the feathers and bone materials

The feather and bone chemistries were further studied via *in-situ* micro-XRF imaging on several sub-areas of interest with higher spatial resolution. The acquisition conditions for the tail measurement where 393 × 193 pixel for a total of 75.8 k pixel/spectra, acquisition time per spectrum was 100 ms and the pixel size 400 μm. The total measurement time was 2h17min. As shown in Fig. 2 and Fig. 3, the distributions of Fe, Ti, Cu, Mn, and Ni match well with the strap pattern of the feathers. The enrichment of many of these elements associated to feather-structures has been also previously reported from synchrotron studies with fossil feathers [1, 6, 12]. Iron level seems relatively lower in the feathers than that in the immediate surrounding remains of body tissue, while Ti, Cu, Mn and Ni seem relatively higher in the feathers than those in the immediate surrounding remains of body tissue (Fig. 2). Notably, unlike the strapped pattern seen on the tail feather (Fig. 2d), the pattern of Cu is not clearly associated with the feathers on the pelvis (Fig. 3d). The Cu and Mn levels apparently vary on feathers of different body parts. The tail feathers have richer Cu than the pelvis feathers, while the Mn level is higher in the pelvic feathers than in the tail feathers (Tables S1-S3). However, these variations in feather samples are not detected for Fe, Ti and Ni. Even though Zn, as well as Ca, Fe, Cu and Mn are known to be typically associated with melanin pigmentation [1, 6, 11, 31, 32], we do not find a connection between Zn and the feathers on the specimen DLXH 1218. Cu and Ni are also distributed in the bone materials, while Fe, Ti and Mn are not significantly enriched in the bones (Figs. 2, 3).

**Fig. 3.**
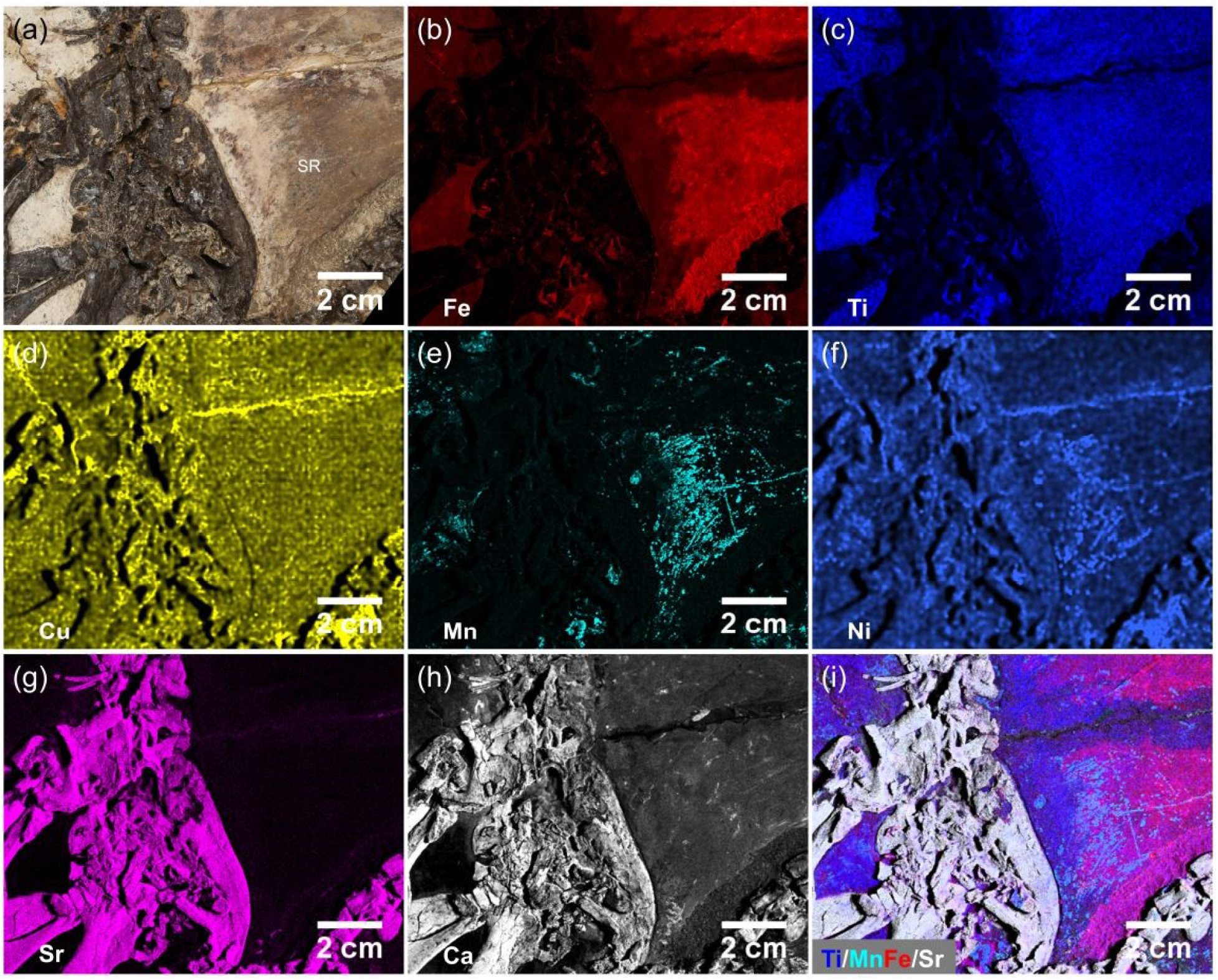
(a) Light photo of a small region of the *Jianianhualong* hip. The bone materials (represented by the black color) and soft tissue (SR) remnants including the feathers can be observed based on light photo imaging. (b)-(n) Micro-XRF detail maps of Fe (b), Ti (c), Cu (d), Mn (e), Ni (f), Sr (g), Ca (h) distribution within bone materials and feathers of the *Jianianhualong* hip. (i) Combined intensity map of four elements. Blue, Ti; aqua, Mn; red, Fe; white, Sr.

Besides Ca, P and S, the heavy elements of Sr, Th, Ce and Y are strongly associated with the skeleton (Figs. 1-3, Tables S1-S4). Comparatively, Sr and Ce levels seem heavily associated with the skeleton and are relatively low in other regions including the slab matrix and the soft tissue remains, as mentioned above.

### 3.3 Sr- and Ca-XRF imaging the morphology and preservation of the skeleton

Both Ca and Sr are accumulated in the bone materials, but Ca is also observed in the soft tissue remains and the slab. As a contrast, Sr seems to be heavily and uniformly accumulated with the skeleton, but present at a very low level in other regions (Figs. 1-3). The Sr/Ca ratio is generally consistent in most part of the skeleton (Tables S1-S4). Fig. 4 shows a comparison of XRF maps of Ca and Sr for the cranium, the pes and the tibiotarsus. Though no novel morphologies are revealed via the chemical imaging, it is clear that, compared to the Ca-maps, the Sr-maps significantly improve the skeletal image by having a better contrast and making the sutures and outlines of bones more visible. Furthermore, comparisons between the Ca and Sr elemental maps may also provide insights into the taphonomy and the process of the fossilization.

**Fig. 4.**
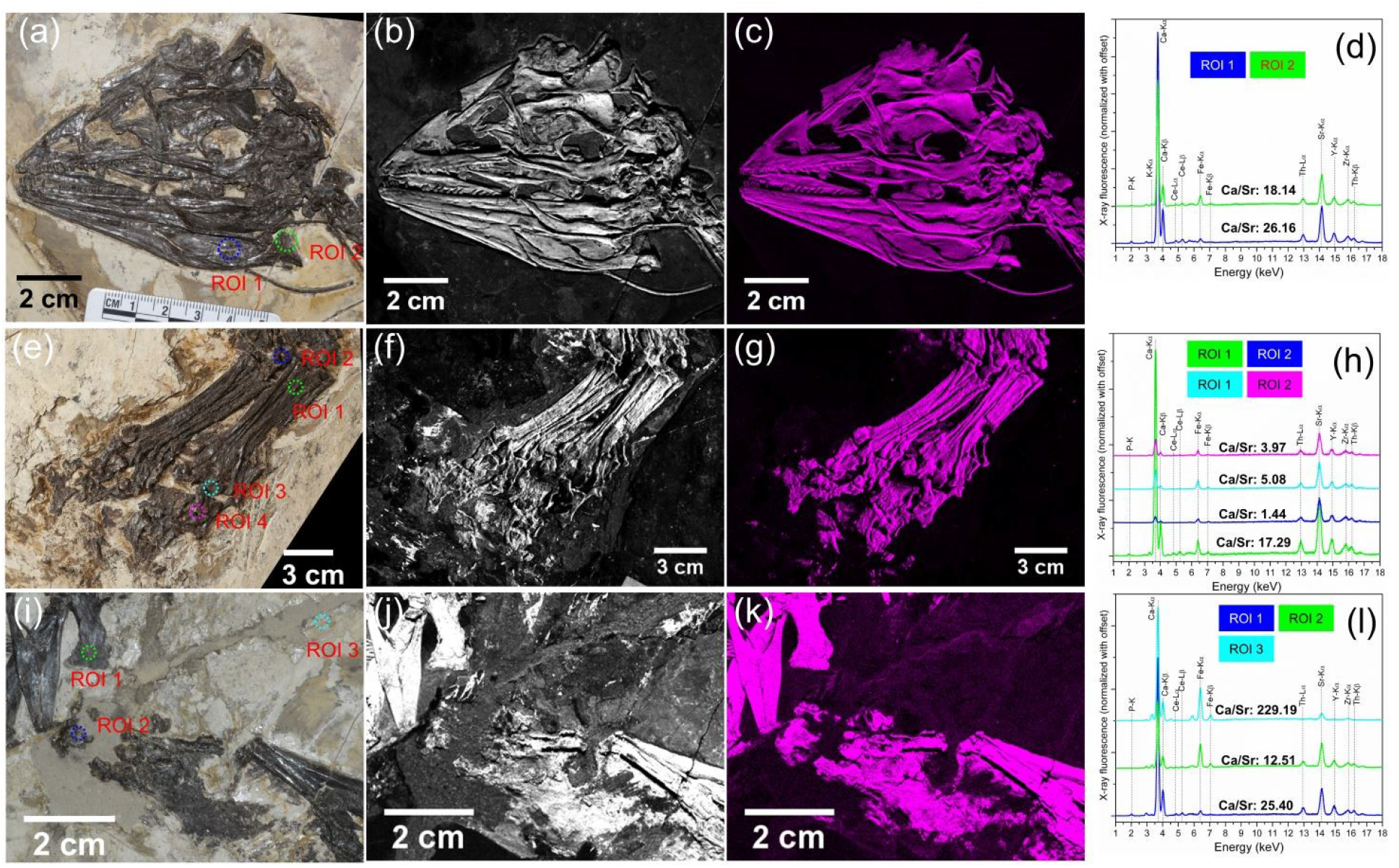
Micro-XRF detail maps of Ca and Sr distribution within the cranium (a)-(c), the pes (e)-(g), and the tibiotarsus (i)-(k) of the *Jianianhualong.* The first column indicates the corresponding light photos, the second for Ca distribution maps, the third for Sr distribution maps. The fourth column indicates the XRF spectrum originated from a small region of interest (ROI) within the cranium (d), the pes (h), and the tibiotarsus (l). Notably, several small regions of interest (ROI) are circled out within the light photos of the cranium (a), the pes (e), and the tibiotarsus (i) of the *Jianianhualong*, and their corresponding XRF spectra are comparatively shown in the panels (d), (h), and (l), respectively.

For instance, the pedal digits, the right metatarsus and the right calcaneum show distinct calcium removal compared to the rest parts of the skeleton, while the strontium level remains stable, which may be caused by the damage of the bone that makes calcium easier to be carried away during fossilization (Fig. 4). Similarly, a calcium removal is present ventral to the distaldorsal process of the right ischium possibly due to a breakage, while the Sr level is relatively stable and the strontium map outlines the shape of the distaldorsal process of the ischium correctly (Fig. 4).

Ambiguously shaped dark colored matters are preserved near the hindlimbs of the specimen DLXH 1218, close to the proximal portion of the tibiotarsus and the distal region of the pes (Fig. 4, Table S3). A high Sr level and high Sr/Ca ratio detected match the shape of the dark colored regions, reaffirming these are remains of smashed bone materials rather than imprints of feathers or other non-skeletal structures, especially considering the proximal portion of the tibiotarsus is severely fragmented and the pedal digits are damaged, which makes the bone materials easily scattered around the damaged bone.

It should be pointed out that a careful examination of the specimen does not reveal new morphologies that have not been already observed under normal light. No trace of cartilage remains are detected associated with the sternum based on Ca, Sr or other elemental maps (Fig. S4), which supports the hypothesis the sternal plates are absent in troodontids [33].

### 3.4 Structure and chemistry of the claw sheath

Interestingly, the claw sheaths showed elemental signatures unnoted previously via chemical imaging (Fig. 5). The sheaths are preserved surrounding the distal portion of manual and pedal distal phalanges in the specimen DLXH 1218. The sheaths have elevated Ca, P and Sr levels, but have very low K, Fe and Mn levels, which matches the pattern of other bones (Fig. 5a, Table S3). A high P profile of the sheaths was also noticed in the Thermopolis *Archaeopteryx, Shuvuuia* and *Citipati* [12, 34, 35]. The Ca level is relatively low as a result of richer Ca in the matrix of Thermopolis *Archaeopteryx* [12], yet a high Ca level is present in the sheaths of *Citipati* and *Shuvuuia* [12, 34, 35] as in *Jianianhualong,* indicating preservation as calcium phosphate. The S level is higher in the sheaths than in the matrix in DLXH 1218 as in the Thermopolis *Archaeopteryx* and *Shuvuuia* [12, 34]. Furthermore, the previous study [12] was unable to distinguish the sheath from the bony part of the claw with elemental maps, but the sheath of DLXH 1218 shows a dorsal rim (unguis) that exhibits high levels of Sr, Th, Ce and Y similar to that in the bone, and these elements are significantly removed from the ventral portion of the sheath blade. Even though the unguis and the sheath blade were both originally keratinous, the chemical distribution in the claw sheath may indicate that the unguis and the sheath blade have compositional or ultrastructural differences in life.

**Fig. 5.**
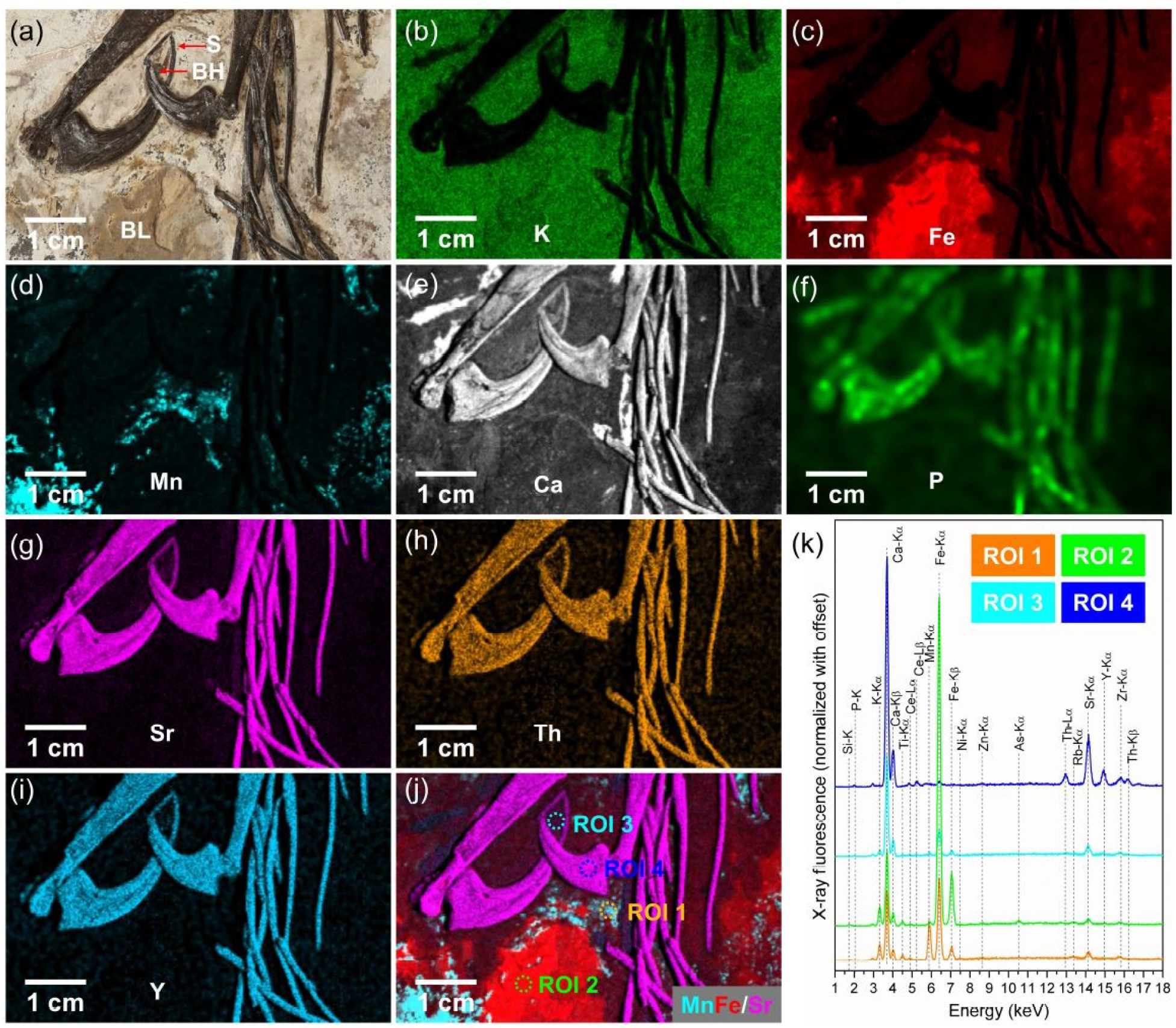
(a) Light photo of a small region of the *Jianianhualong* claw. The bone materials of the claw (C) (represented by the black color), the corresponding sheaths (S) and horn (H) can be observed based on light photo imaging. (b)-(j) Micro-XRF detail maps of K (b), Fe (c), Mn (d), Ca (e), P (f), Sr (g), Th (h), and Y (i) distribution within the bone and sheath materials of the *Jianianhualong* claw. (j) Combined intensity map of three elements. Aqua, Mn; red, Fe; magenta, Sr. (k) XRF spectra originated from the four small regions of interest (ROI) circled out by different colors in the panel (j).

## 4. Discussion and conclusions

### 4.1 Information revealed by micro-XRF chemical imaging

Applying large-area micro-XRF on DLXH 1218 reveals the surface elemental distribution on the biological remains of the specimen and its surrounding geological matrix. Unlike chemical mappings in previous studies that focus on the lighter elements [12, 29], the micro-XRF imaging is also able to detect the distribution of heavier elements. Generally, the distributions of Fe, Ti, Cu, Mn and Ni are associated with the soft remains, and they match well with the pattern of the feathers. Heavier elements Sr, Th, Y and Ce are strongly associated with the skeleton. Though no novel morphology was uncovered by the chemical mapping, compositional variations are noted in some structures to reflect either biological or taphonomic features. For instance, the Sr variation among the unguis, the sheath blade and the bony claw likely indicate the compositional differences among these structures in life. The Ca variation on some parts of the skeleton (e.g. between pedal digits and the metatarsus) possibly showed the Ca deleting during fossilization.

Combining chemical information of the lighter and heavier elements can provide insights to the material exchange of the fossil remains and its surrounding rocks during fossilization. Notably, a crack-like structure posterior to the proximal caudal showed interesting elemental abnormal (Fig. 1). The Ca, P and S levels in this crack are similar to that of the skeleton, while the Sr, Th, Y and Ce levels are very low in this crack compared with the bones. Unlike other cracks that are filled with glue derived from the fossil preparation, the As, Cu, Ni, Ga and Zn levels are also low in this crack (Fig. 1, Fig. S2). And therefore, we infer this crack was formed during the fossilization, and the water flux carried the lighter elements Ca, P and S from the animal remains into this crack. On the other hand, heavier elements like Sr, Th, Y and Ce were more easily accumulated in the bone materials and were not carried away by water flux, which may further indicate at least part of these heavier elements in the skeleton are original, as supported by the fact that Sr participates significantly in bioapatite mineralization throughout the life of an animal [36].

### 4.2 Comparisons with chemistry of other fossil materials

Comparisons of chemical mappings of DLXH 1218 and other vertebrate fossils not only show consistent patterns of chemical accumulation in bones and feathers, but also indicate elements behave differently depending on the chemical composition of surrounding rocks, as previously noticed [8, 30, 37]. Rossi and co-workers [14] concluded that elemental signature of vertebrate fossils is tissue-specific and is controlled by many biological and environmental factors, which is congruent with our findings in DLXH 1218 that both similarities and differences could be recognized in the elemental distributions on feathers and bones compared with other vertebrate fossils. As the composition of the bones is representative for the depositional environment and most likely local condition of fossilization, this variation of local compositional fingerprint could be used to assign poorly or not correct documented fossil to certain location. Here for additional systematic work is needed.

Specifically, S, P and Ca are the most commonly examined elements in chemical studies based on fossil materials [30, 38]. Consistently, micro-XRF with DLXH 1218 showed that S, P and Ca are associated with the organic matter (both the skeleton and soft tissues), while Ca levels are slightly higher in the skeleton, a pattern similar to previously examined fossil specimens [1, 12, 31]. Comparatively, the Ca level in the matrix of DLXH 1218 is relatively low, different from the Thermopolis *Archaeopteryx*, where the matrix rock is calcium-rich [12]. The claw sheaths preserved in DLXH 1218 have elevated Ca and P levels, congruent with the patterns noticed in the Thermopolis *Archaeopteryx*,*Shuvuuia* and *Citipati* specimens [12, 34, 35].

Some elements such as Ca, Zn, Mn, Cu and Ni have been noticed to be associated with melanin pigmentation in modern and fossil feathers [1, 6, 11, 18, 29, 31, 32]. In the fossils of *Jianianhualong*, *Anchiornis*, *Archaeopteryx*, *Gansus*, and other extinct and living birds, the Cu level usually varies on different feather samples of an individual and is generally higher in the feathers than in the bones [6, 12, 29, 31]. Though in DLXH 1218, the distributions of Ca, Cu, Mn and Ni match the plumage shapes, no obvious enrichment of Zn has been observed with either feathers or bones, which is also different from the Thermopolis *Archaeopteryx*, *Confuciusornis* and the fossil coliiformes bird (AMNH FARB 30806) where the feathers and bone materials have elevated Zn level [6, 12, 29]. Instead, the highest level of Zn is present in the glue added in DLXH 1218. In DLXH 1218, aside from the matching pattern of Mn and the pelvic feathers, high Mn level areas are also scattered on the sediments, similar to the Thermopolis *Archaeopteryx* and the Green River feathers (BHI-6358, BHI-6319), where the Mn level is generally higher in the matrix than in the feathers [12, 31], but unlike that in *Confuciusornis, Gansus* and the fossil coliiformes specimens, where the Mn is rich in feathers but depleted from parts of the matrix [6, 29, 31]. A high Ti level is also associated with the feathers preserved in DLXH 1218, a pattern that has not been reported in other fossil materials, although Ti level variation has been detected between two *Gansus* feathers and as a contrast Ti is absent in some modern bird feathers [31].

Though a high Fe level is also known in some melanized structures, fossil feathers preserved in DLXH 1218 show a relatively lower Fe level than in the surrounding areas, similar to that in the Thermopolis *Archaeopteryx* [12]. DLXH 1218 resembles the coliiformes bird in having a very low Fe level in bones, while the bone of the Thermopolis *Archaeopteryx* is more iron-rich than its feathers. Barden and co-workers [31], noted that the feathers of the modern marabou stork and white-naped crane did not contain noticeable Fe with EDS (energy dispersive X-ray spectrometry) analyses, and the Fe level was low and varied in two fossilized *Gansus* feathers. We suspect that the plumage iron level is somewhat influenced by the matrix.

Strontium is strongly associated in the bones preserved in DLXH 1218, which is not surprising as Sr is commonly participated in forming bone bioapatite, and relatively high Sr levels have been found in many fossil bone materials [30, 36, 37, 39]. Different chemical behaviors of Sr and Y in calcium phosphates and carbonates have been suggested by Gueriau and co-workers [30, 39] based on fish and non-vertebrate fossils, yet the distribution of Sr and Y are congruent in DLXH 1218, as the bones are calcium phosphates only.

In summary, this study performed a large-area Micro-XRF scanning on the holotype of *Jianianhualong tengi*, and obtained for the first time the chemical maps of the whole specimen with resolution at the micron scale. A detailed analysis of elemental distribution and compositional variations in the skeleton, feather and the surrounding matrix demonstrated that the main elements of organisms such as S, P and Ca, as expected, were found to be associated with both the skeleton and soft tissue remains, while Ca was relatively higher in the former than that in the latter. Some elements such as Cu, Mn, Ni and Ti were detected to match the plumage shapes, while other elements such as Sr, Y, and Ce were strongly associated with the skeleton. A careful examination of the chemical signature of the bones and feathers of the *Jianianhualong* DLXH 1218 provide new information on the taphonomy of this important fossil and also paleobiology of this key species for understanding paravian as well as troodontid evolution, and also gave important clues for further study on a few small regions of interest that can be sampled without damage to the specimen.

## Supporting information

supplementary tables and associated figures

## ACKNOWLEDGMENTS

This study was supported financially by the National Natural Science Foundation of China (grants 41890843, 41688103, 41920104009, 41621004 and 41972025), the Laboratory for Marine Geology, Qingdao National Laboratory for Marine Science and Technology (grant MGQNLM201704). We thank Mrs Wang Lixia for inspiring us with this project. We thank Mr. Wu Baojun (UCAS), Mr. Meng Liang (DLTV-4) and Mrs Zhang Jiang (CCTV-10) for their assistant in the organization of the research project. We also thank Mrs Gao Xia and Zhou Jian at the DLXH museum for their assistant in the experiment. We are grateful for Mrs Su Hui and Sun Liqing at the Boyue Instruments (Shanghai) Co., Ltd. for their efficient job in the instrument coordination. J.H.L. benefited from discussions colleagues in Office 442 at IGGCAS.

