## supplementary figures for "Micro-XRF study of the troodontid dinosaur *Jianianhualong tengi* reveals new biological and taphonomical signals"

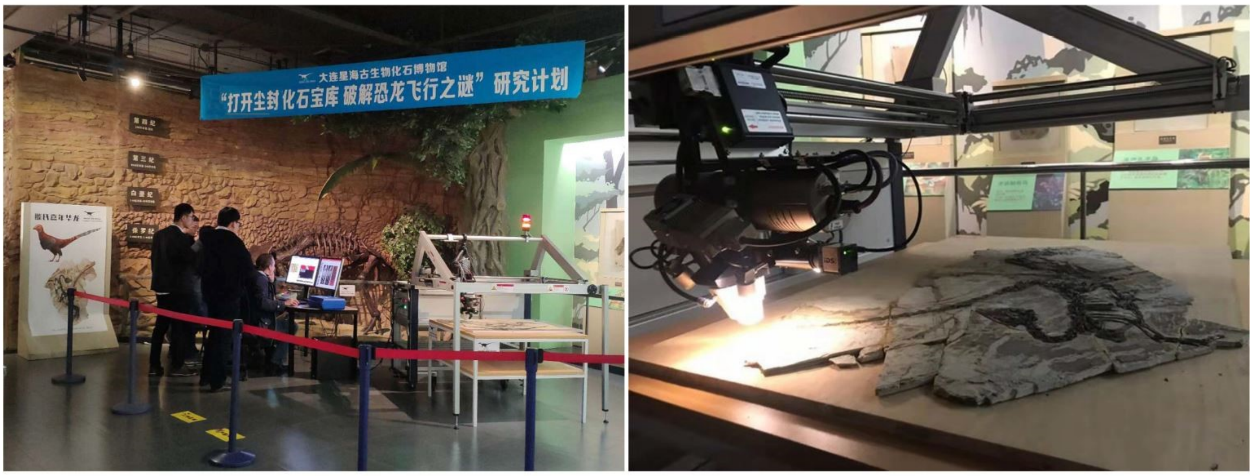


Fig. S1. Micro-XRF (Bruker M6 Jetstream) setup within the Xinghai Paleontological Museum of Dalian used for scanning the *Jianianhualong*.

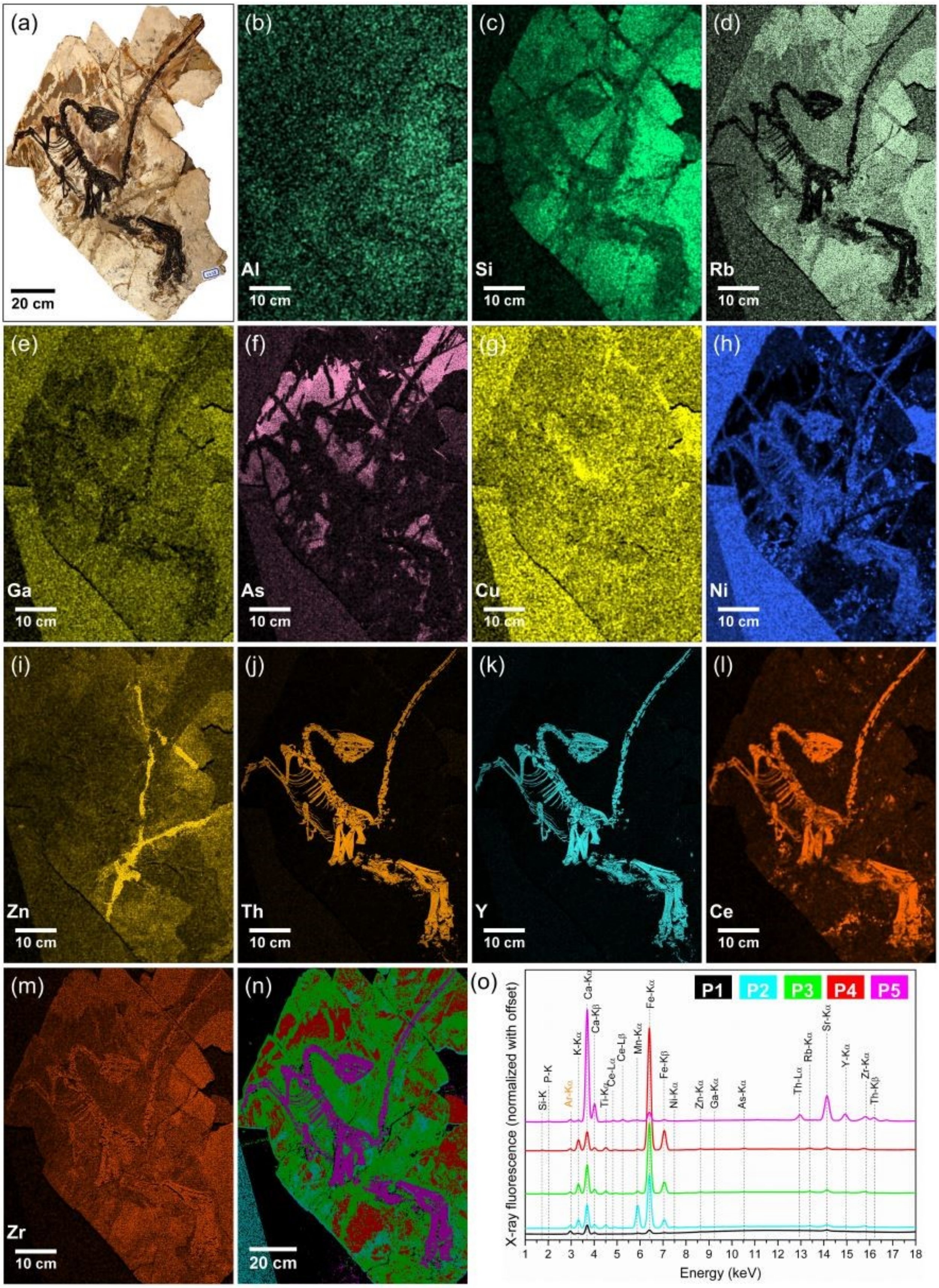
(a)
20 cm
(b)
Al
10 cm
(c)
Si
10 cm
(d)
Rb
10 cm
(e)
Ga
10 cm
(f)
As
10 cm
(g)
Cu
10 cm
(h)
Ni
10 cm
(i)
Zn
10 cm
(j)
Th
10 cm
(k)
Y
10 cm
(l)
Ce
10 cm
(m)
Zr
10 cm
(n)
20 cm
(o)
P1 P2 P3 P4 P5
X-ray fluorescence (normalized with offset)
Energy (keV)
Si-K P-K Ar-Kα K-Kα Ca-Kα Ca-Kβ Ti-Kα Ce-Lα Ce-Lβ Mn-Kα Fe-Kα Fe-Kβ Ni-Kα Zn-Kα Ga-Kα As-Kα Th-Lα Rb-Kα Sr-Kα Y-Kα Zr-Kα Th-Kβ
1 2 3 4 5 6 7 8 9 10 11 12 13 14 15 16 17 18

Fig. S2. (a) Light photo of the *Jianianhualong*. (b)-(m) Intensity maps in the rays of Al-K (b), Si-K (c), Rb-Kα (d), Ga-Kα (e), As-Kα (f), Cu-Kα (g), Ni-Kα (h), Zn-Kα (i), Th-Lα (j), Y-Kα (k), Zr-Kα (l), and Ce-Lβ (m). (n) Reconstructed image with PCA-filtered spectra originated from the support plank, the Mn-rich region, the white layer of the slab rich in K, the brown layer of the slab rich in Fe, and the skeleton. (o) Recalculated XRF spectra for the five phases obtained from the PCA-filtered data. The phase 1 (P1), phase 2 (P2), phase 3 (P3), phase 4 (P4), and phase 5 (P5) match well with the support plank (black color), the Mn-rich region in the slab (aqua color), the white layer of the slab (green color), the brown layer of the slab (red color), and the skeleton of *Jianianhualong* (magenta color), respectively.

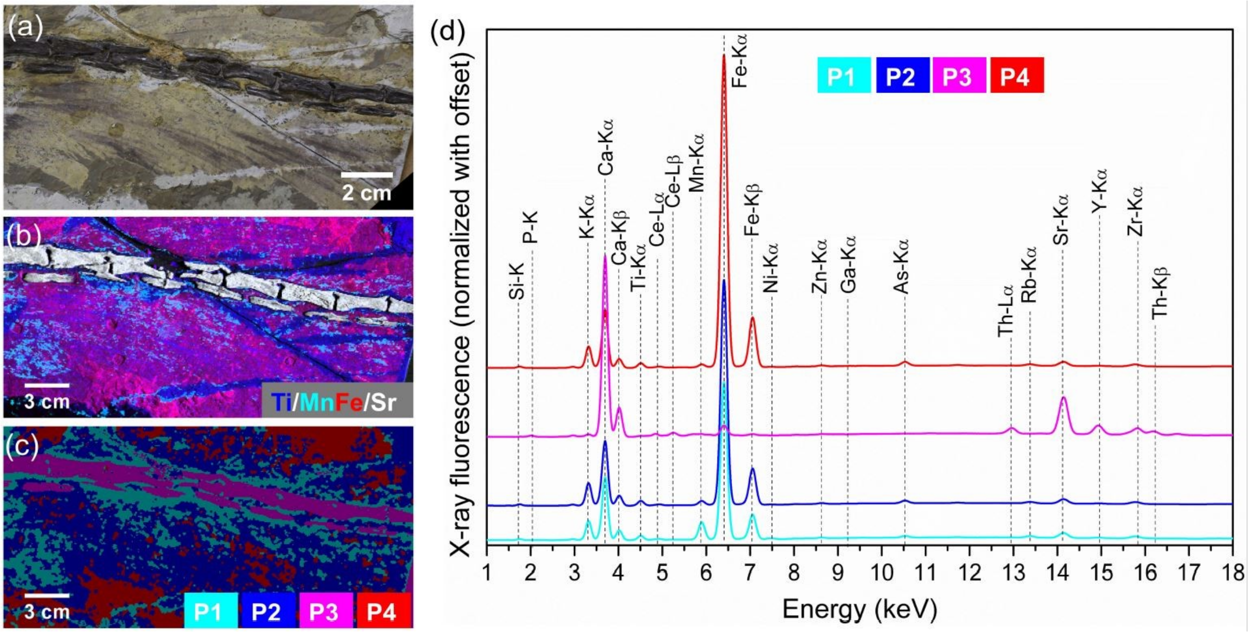


Fig. S3. (a) Light photo of a small region of the *Jianianhualong* tail. (b) Combined XRF intensity map of four elements. Blue, Ti; aqua, Mn; red, Fe; white, Sr. (c) Reconstructed image with PCA-filtered spectra originated from the Mn-rich region, the soft tissue remnants including the feathers, the skeleton, and the brown layer of the slab. (d) Recalculated XRF spectra for the four phases obtained from the PCA-filtered data.

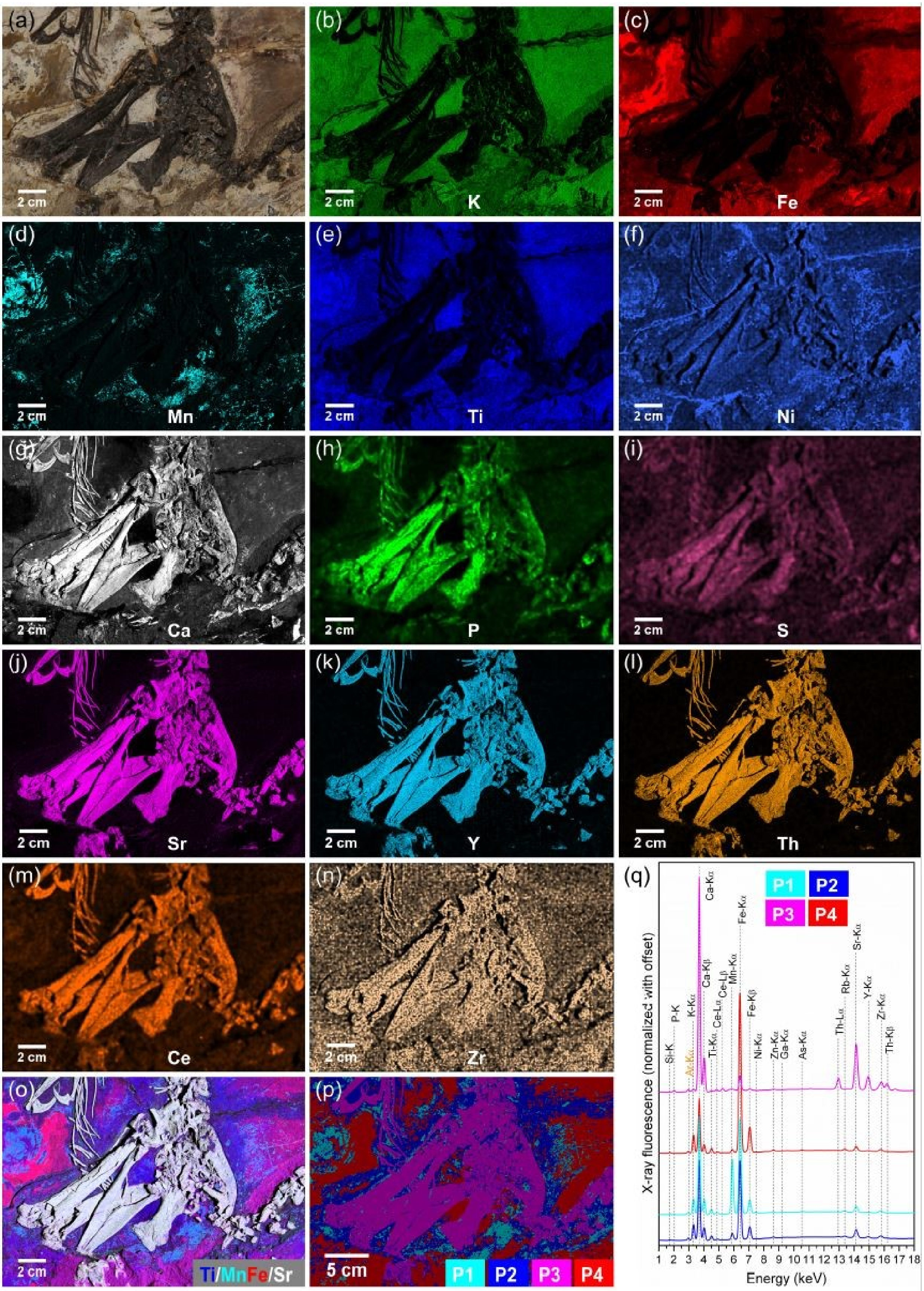

(a)
2 cm
(b)
2 cm
K
(c)
2 cm
Fe
(d)
2 cm
Mn
(e)
2 cm
Ti
(f)
2 cm
Ni
(g)
2 cm
Ca
(h)
2 cm
P
(i)
2 cm
S
(j)
2 cm
Sr
(k)
2 cm
Y
(l)
2 cm
Th
(m)
2 cm
Ce
(n)
2 cm
Zr
(o)
2 cm
Ti/MnFe/Sr
(p)
5 cm
P1 P2 P3 P4
(q)
P1 P2 P3 P4
X-ray fluorescence (normalized with offset)
Energy (keV)
Si-K P-K Ar-Kα K-Kα Ca-Kα Ca-Kβ Ti-Kα Ce-Lα Ce-Lβ Mn-Kα Fe-Kα Fe-Kβ Ni-Kα Zn-Kα Ga-Kα As-Kα Th-Lα Rb-Kα Sr-Kα Y-Kα Zr-Kα Th-Kβ
1 2 3 4 5 6 7 8 9 10 11 12 13 14 15 16 17 18

Fig. S4. (a) Light photo of the *Jianianhualong* hip and claws. (b)-(n) Micro-XRF detail maps of K (b), Fe (c), Mn (d),Ti (e), Ni (f), Ca (g), P (h), S (i), Sr (j), Y (k), Th (l), Zr (m), and Ce (n) distribution within the bone materials and the soft tissue remnants of the *Jianianhualong* hip and claws. (o) Combined XRF intensity map of four elements. Blue, Ti; aqua, Mn; red, Fe; white, Sr. (p) Reconstructed image with PCA-filtered spectra originated from the Mn-rich region, the soft tissue remnants including the feathers, the skeleton, and the brown layer of the slab. (q) Recalculated XRF spectra for the four phases obtained from the PCA-filtered data.

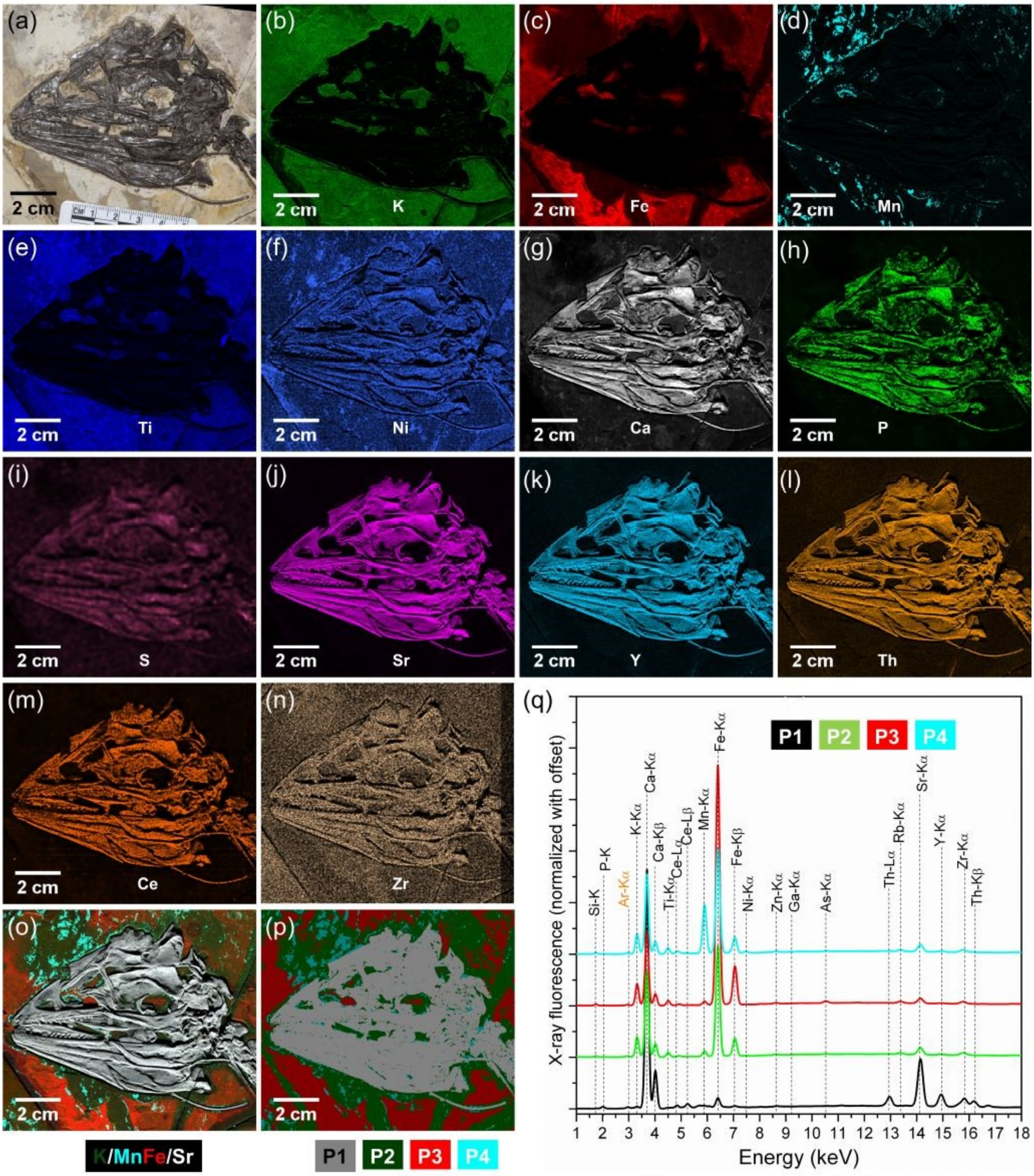


Fig. S5. (a) Light photo of the *Jianianhualong* cranium. (b)-(n) Micro-XRF detail maps of K (b), Fe (c), Mn (d),Ti (e), Ni (f), Ca (g), P (h), S (i), Sr (j), Y (k), Th (l), Zr (m), and Ce (n) distribution within the bone materials and the soft tissue remnants of the *Jianianhualong* cranium. (o) Combined XRF intensity map of four elements. Blue, Ti; aqua, Mn; red, Fe; white, Sr. (p) Reconstructed image with PCA-filtered spectra originated from the skeleton, the white layer of the slab, the brown layer of the slab, and the Mn-rich region. (q) Recalculated XRF spectra for the four phases obtained from the PCA-filtered data.

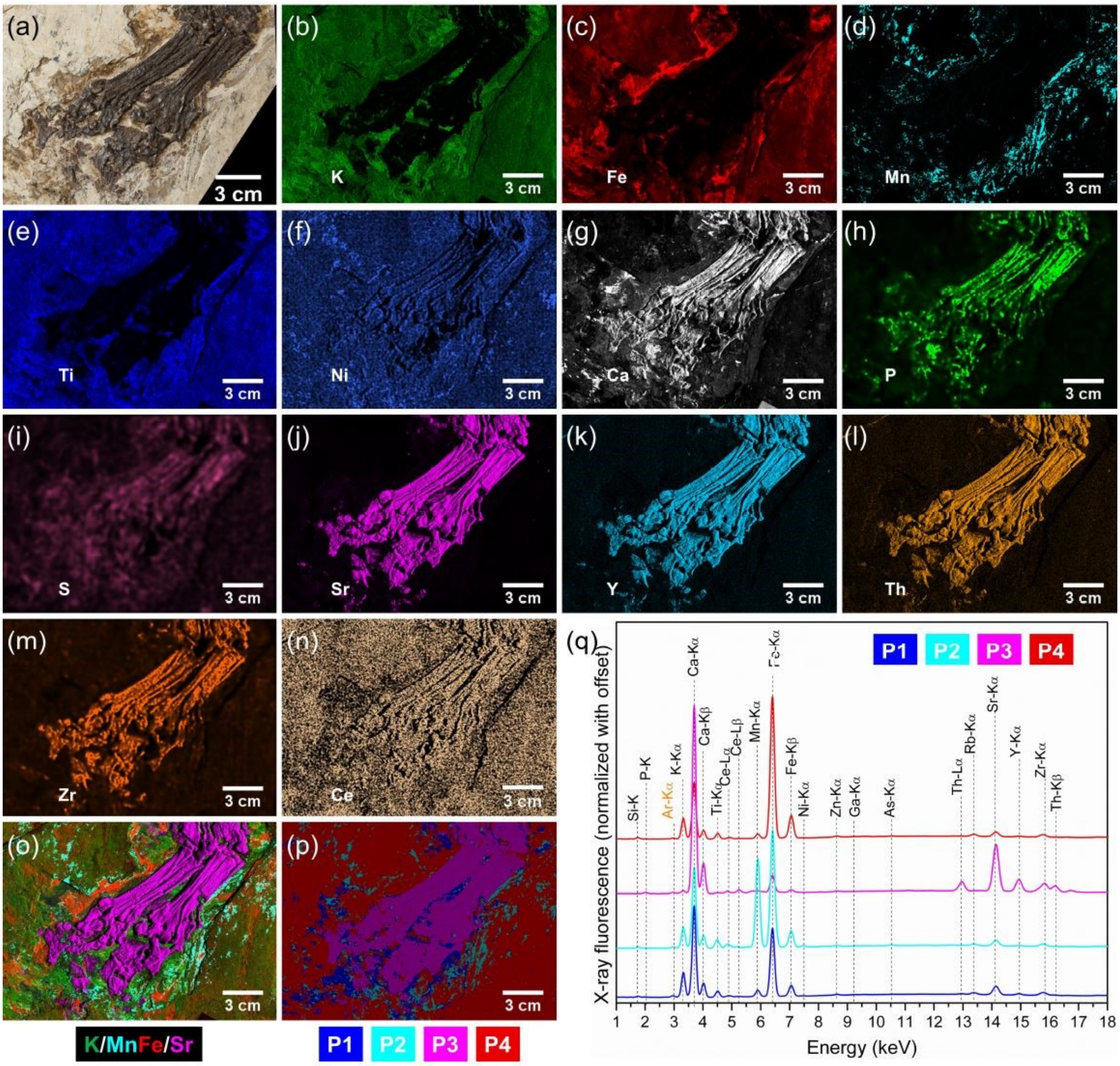


Fig. S6. (a) Light photo of the *Jianianhualong* pes. (b)-(n) Micro-XRF detail maps of K (b), Fe (c), Mn (d),Ti (e), Ni (f), Ca (g), P (h), S (i), Sr (j), Y (k), Th (l), Zr (m), and Ce (n) distribution within the bone materials and the soft tissue remnants of the *Jianianhualong* pes. (o) Combined XRF intensity map of four elements. Green, K; aqua, Mn; red, Fe; magenta, Sr. (p) Reconstructed image with PCA-filtered spectra originated from the soft tissue remnants, the Mn-rich region, the skeleton, and the white layer of the slab. (q) Recalculated XRF spectra for the four phases obtained from the PCA-filtered data.
