## supplementary tables and associated figures for "Micro-XRF study of the troodontid dinosaur *Jianianhualong tengi* reveals new biological and taphonomical signals"

Table S1 Comparison of element concentration in different regions of interest (ROI) on the *Jianianhualong* fossil in net intensity (cps)

| ROI | Ca | Si | Fe | K | Al | P | Ti | Mn | Sr | Th | Ce | Y | Zr | Rb | Zn | As | S | Cu | Ga | Ni |
|---|---|---|---|---|---|---|---|---|---|---|---|---|---|---|---|---|---|---|---|---|
| Overall | 4629 | 37 | 10509 | 951 | 1 | 10 | 340 | 526 | 1050 | 200 | 62 | 188 | 159 | 178 | 110 | 78 | - | 25 | 31 | 11 |
| WL-1 | 2050 | 39 | 8674 | 1190 | - | 0.3 | 441 | 117 | 403 | - | - | - | 227 | 220 | 72 | - | - | - | 19 | - |
| WL-2 | 1150 | 36 | 9535 | 1291 | - | - | 491 | 75 | 393 | - | - | - | 238 | 551 | 111 | 11 | - | 5 | 15 | - |
| BL-3 | 4214 | 22 | 21961 | 767 | - | - | 266 | 198 | 493 | - | - | - | 86 | 108 | 45 | 338 | - | 28 | - | - |
| BL-4 | 4283 | 28 | 27201 | 1051 | - | 0.4 | 359 | 197 | 557 | - | 14 | - | 125 | 154 | 73 | 428 | - | 27 | - | - |
| OL-5 | 4795 | 25 | 18077 | 1198 | - | - | 321 | 223 | 604 | - | 3 | - | 113 | 137 | 55 | 314 | - | 17 | 4 | - |
| OL-6 | 5680 | 28 | 21898 | 1298 | - | - | 369 | 234 | 626 | - | - | - | 97 | 117 | 41 | 284 | - | 24 | - | - |
| FL-7 | 8519 | 78 | 22185 | 1962 | 1 | - | 575 | 1526 | 786 | - | 4 | 3 | 190 | 197 | 85 | 236 | - | 71 | 12 | 23 |
| FL-8 | 5950 | 32 | 29997 | 1764 | - | - | 563 | 270 | 579 | - | - | - | 94 | 93 | 42 | 460 | - | 43 | - | - |
| FL-9 | 4018 | 39 | 27978 | 1508 | - | - | 423 | 179 | 447 | - | - | - | 122 | 125 | 46 | 449 | - | 33 | - | - |
| SR-10 | 20405 | - | 659 | 18 | - | 93 | 20 | 90 | 5836 | 970 | 496 | 1365 | - | - | 53 | 8 | - | - | - | - |
| SR-11 | 27815 | - | 939 | 53 | - | 205 | 44 | 110 | 7973 | 1690 | 487 | 2144 | - | - | 63 | - | - | - | - | - |
| SR-12 | 17238 | 4 | 2392 | 121 | - | 67 | 47 | 212 | 6516 | 974 | 376 | 1623 | - | - | 78 | - | - | 12 | - | - |
| RR-13 | 6688 | 86 | 24408 | 1466 | - | - | 651 | 476 | 910 | - | 10 | 7 | 228 | 197 | 585 | 35 | - | 14 | 12 | - |
| RR-14 | 6500 | 77 | 26136 | 1452 | - | 4 | 667 | 405 | 717 | - | 5 | - | 245 | 227 | 498 | 30 | - | 15 | - | - |

Notes: WL: white layer, BL: brawn layer, OL: organic layer, FL: feather layer, SR: skeleton region, RR: restoration region. An increasing of the relative concentration compared to its overall is showing by red color, while a decreasing by green color.

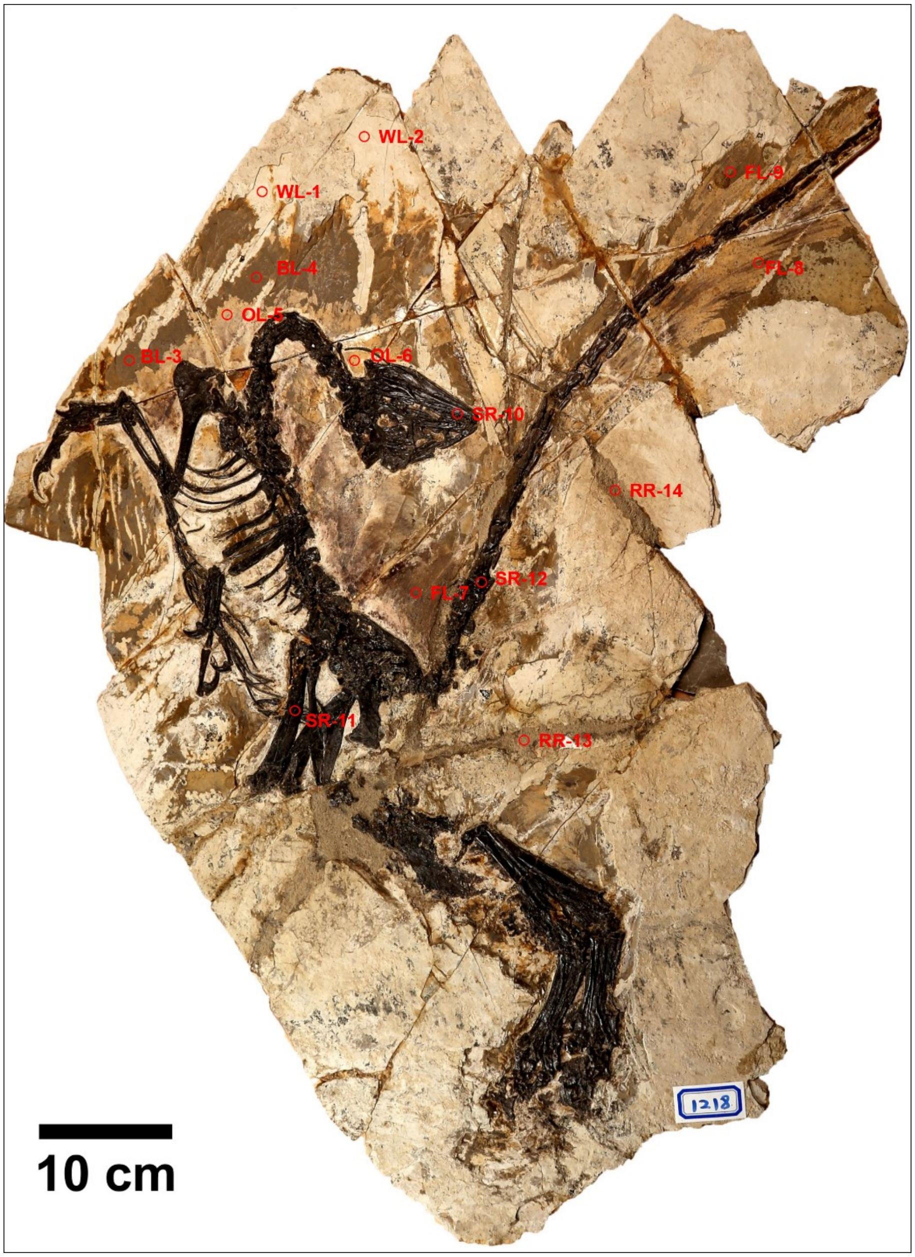
WL-2
WL-1
BL-4
OL-5
BL-3
OL-6
FL-9
FL-8
SR-10
RR-14
SR-12
FL-7
SR-11
RR-13
1218
10 cm

Table S2 Comparison of element concentration in different regions of interest (ROI) on the tail part of the *Jianianhualong* fossil in net intensity (cps)

| ROI | Ca | Si | Fe | K | P | Ti | Mn | Sr | Th | Ce | Y | Zr | Rb | Zn | As | S | Cu | Ga | Ni |
|---|---|---|---|---|---|---|---|---|---|---|---|---|---|---|---|---|---|---|---|
| Overall | 8421 | 106 | 27205 | 1974 | 14 | 517 | 792 | 1494 | 206 | 100 | 219 | 233 | 243 | 139 | 389 | 2 | 42 | 26 | 30 |
| SL-1 | 25487 | 1 | 487 | - | 222 | 42 | 116 | 7544 | 990 | 590 | 1633 | - | - | 37 | - | - | 6 | - | - |
| SL-2 | 20990 | - | 971 | 117 | 73 | 45 | 119 | 7557 | 1025 | 572 | 1721 | - | - | 51 | - | - | - | - | - |
| OL-3 | 5375 | 154 | 39593 | 2580 | - | 574 | 943 | 729 | - | - | 7 | 222 | 182 | 148 | 622 | - | 27 | 16 | 12 |
| OL-4 | 6810 | 143 | 39005 | 2661 | - | 581 | 715 | 683 | - | - | - | 211 | 243 | 101 | 570 | - | 13 | - | - |
| FL-5 | 8002 | 133 | 36609 | 2353 | - | 727 | 383 | 796 | - | 4 | - | 220 | 204 | 91 | 571 | - | 75 | 2 | - |
| FL-6 | 9243 | 118 | 34388 | 2261 | - | 658 | 341 | 773 | - | - | - | 202 | 243 | 93 | 549 | 2 | 91 | - | - |

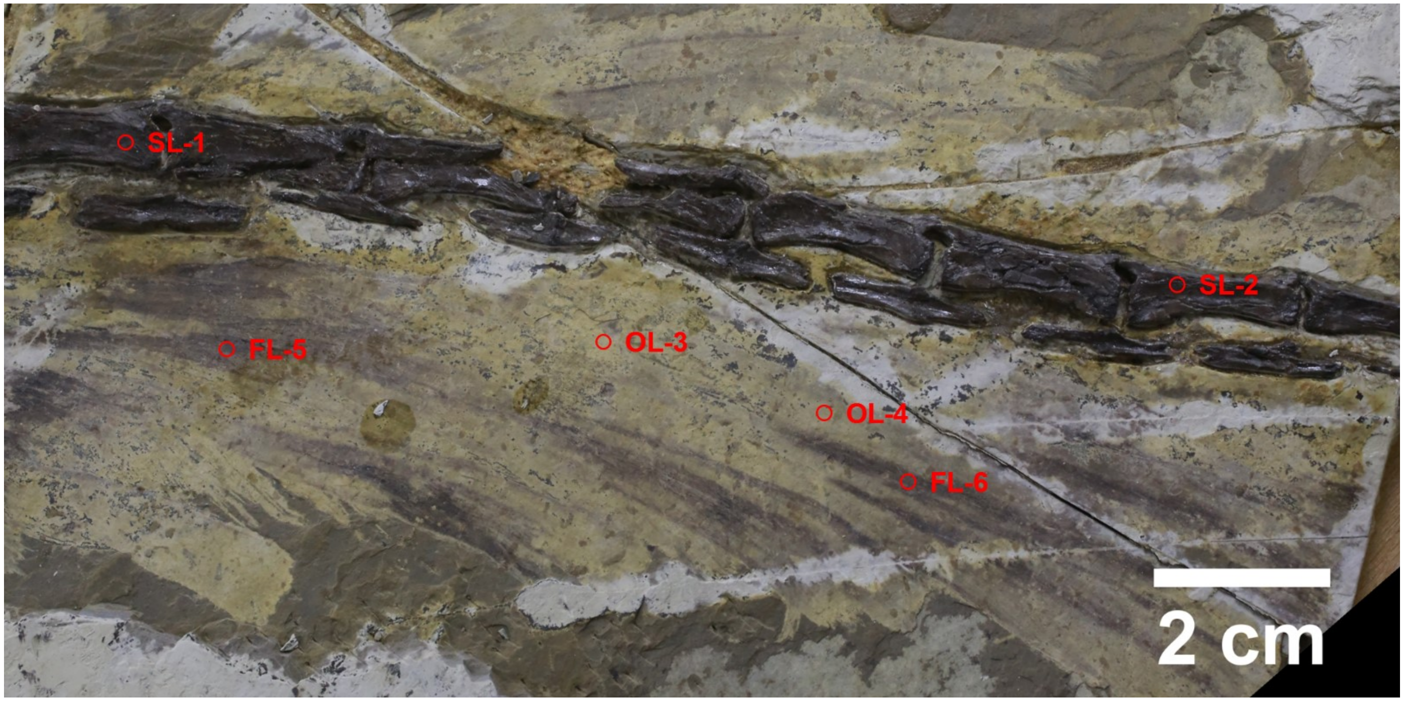

Table S3 Comparison of element concentration in different regions of interest (ROI) on the *Jianianhualong* hip and claws in net intensity (cps)

| ROI | Ca | Si | Fe | K | P | Ti | Mn | Sr | Th | Ce | Y | Zr | Rb | Zn | As | S | Cu | Ga | Ni |
|---|---|---|---|---|---|---|---|---|---|---|---|---|---|---|---|---|---|---|---|
| Overall | 10948 | 49 | 10428 | 1095 | 39 | 391 | 944 | 2818 | 496 | 173 | 671 | 168 | 165 | 155 | 63 | - | 31 | 26 | 24 |
| OL-1 | 6131 | 114 | 11386 | 2247 | - | 769 | 799 | 748 | - | - | - | 234 | 182 | 131 | - | - | - | 9 | 65 |
| FL-2 | 10737 | 76 | 19032 | 1939 | - | 571 | 2700 | 841 | - | - | - | 176 | 205 | 81 | 217 | - | 41 | 2 | 27 |
| SL-3 | 30062 | - | 523 | - | 277 | 36 | 172 | 8084 | 1594 | 592 | 2104 | - | - | 54 | - | - | - | - | - |
| SL-4 | 9186 | 11 | 4487 | 387 | 14 | 118 | 222 | 5396 | 1033 | 115 | 1424 | 24 | - | 117 | - | - | - | - | - |
| CR-5 | 24338 | - | 928 | - | 110 | 30 | 105 | 7071 | 1212 | 582 | 1646 | - | - | 74 | - | - | - | - | - |
| CR-6 | 20103 | 17 | 1155 | 89 | 18 | 4 | 81 | 4902 | 1117 | 453 | 1138 | - | - | 78 | - | - | - | - | - |

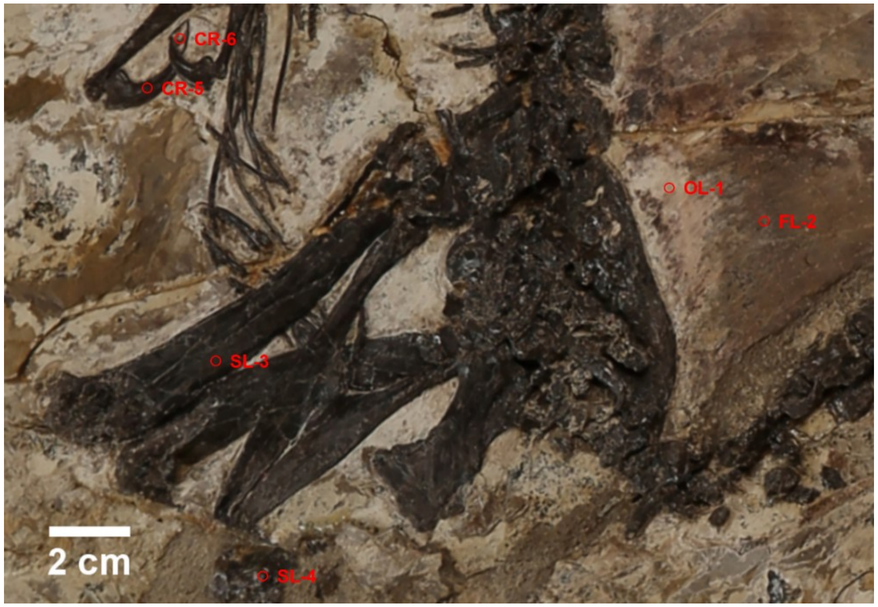

Table S4 Comparison of element concentration in different regions of interest (ROI) on the *Jianianhualong* pes in norm. stoich. C [wt. %

| ROI | Ca | Si | Fe | K | P | Ti | Mn | Sr | Th | Ce | Y | Zr | Rb | Zn | As | S | Cu | Ga | Ni |
|---|---|---|---|---|---|---|---|---|---|---|---|---|---|---|---|---|---|---|---|
| Overall | 8650 | 92 | 12756 | 1565 | 28 | 524 | 1053 | 2215 | 344 | 157 | 451 | 254 | 251 | 139 | 47 | 1 | 22 | 32 | 24 |
| WL-1 | 16447 | 113 | 8032 | 1551 | - | 400 | 658 | 1286 | - | - | - | 80 | - | 33 | - | - | - | - | - |
| BL-2 | 7426 | 101 | 37999 | 1972 | - | 530 | 229 | 795 | - | - | - | 160 | 133 | 21 | 107 | - | 15 | - | - |
| OL-3 | 6862 | 78 | 19356 | 2004 | - | 735 | 5249 | 726 | - | 51 | - | 166 | 149 | 69 | 27 | - | - | - | - |
| SL-4 | 22576 | - | 986 | - | 198 | - | 81 | 7682 | 1422 | 392 | 1930 | - | - | 39 | - | - | - | - | - |
| SL-5 | 7267 | - | 3245 | - | 43 | 24 | 20 | 6630 | 1083 | 96 | 1421 | 62 | - | 18 | - | - | - | 8 | - |
| SL-6 | 13285 | 4 | 1452 | 155 | 48 | 39 | 55 | 5499 | 899 | 259 | 1223 | - | - | 132 | - | - | - | - | - |

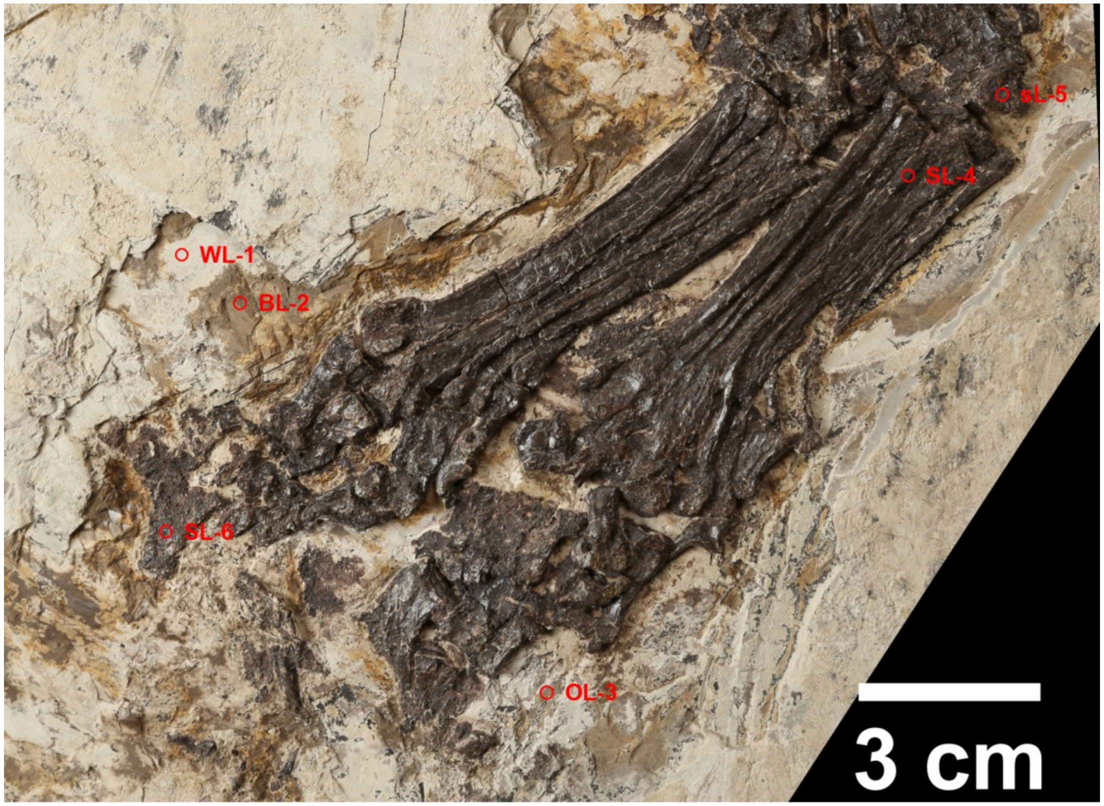
